# Label-free imaging for quality control of cardiomyocyte differentiation

**DOI:** 10.1101/2021.06.01.446615

**Authors:** Tongcheng Qian, Tiffany M. Heaster, Angela R. Houghtaling, Kexin Sun, Kayvan Samimi, Melissa C. Skala

## Abstract

Human pluripotent stem cell (hPSC)-derived cardiomyocytes provide a promising regenerative cell therapy for cardiovascular patients and an important model system to accelerate drug discovery. However, cost-effective and time-efficient platforms must be developed to evaluate the quality of hPSC-derived cardiomyocytes during biomanufacturing. Here, we developed a non-invasive label-free live cell imaging platform to predict the efficiency of hPSC differentiation into cardiomyocytes. Autofluorescence imaging of metabolic co-enzymes was performed under varying differentiation conditions (cell density, concentration of Wnt signaling activator) across three hPSC lines. Live cell autofluorescence imaging and multivariate classification models provided high accuracy to separate low (< 50%) and high (≥ 50%) differentiation efficiency groups (quantified by cTnT expression on day 12) within 1 day after initiating differentiation (area under the receiver operating characteristic curve, 0.98). This non-invasive and label-free method could be used to avoid batch-to-batch and line-to-line variability in cell manufacturing from hPSCs.

## Introduction

Despite advances in treatment, cardiovascular disease is the leading cause of death worldwide^1^. Globally, about 12% of adults are diagnosed with cardiovascular disease and over 30% of all deaths are caused by cardiovascular disease^1^. The excessive demand of heart transplantation has outpaced the limited number of healthy and functional heart donors^2^. Cell-based regenerative therapy provides a promising treatment for patients suffering from cardiac tissue injury^3, 4^. However, cardiomyocytes (CMs) are terminally differentiated cells with no regenerative capacity^5^. Hence, cost-effective and time-efficient platforms to generate functional CMs with high quality has emerged as an urgent need for cardiac medicine in drug screening, toxicity testing, disease modeling, and regenerative cell therapy.

Human pluripotent stem cells (hPSCs) can differentiate into cells from all three germ layers^6–8^. A variety of methods have been established to generate CMs from hPSCs^9–11^. These hPSC-derived CMs exhibit similar functional phenotypes to their *in vivo* counterparts^11^, including self-contractility and action potentials. hPSC-derived CMs have been used in disease modeling^12, 13^ and drug screening^14^, and hold great potential for regenerative medicine^15, 16^. However, batch-to-batch and line-to-line variability in the differentiation process of hPSCs into CMs has impeded the scale-up of CM manufacturing^17^. For safety, the quality of clinical-graded hPSC-derived CMs must be rigorously evaluated before they can be used for regenerative cell therapy in patients^18^. Current approaches to quantify CM differentiation rely on low-throughput, labor-intensive, and destructive immunofluorescence labelling and electrophysiological measurements^11^. New technologies that can non-invasively monitor CM differentiation in real time and evaluate the differentiation outcome at early stages are needed to effectively optimize biomanufacturing of CMs from stem cells.

Previous studies indicate that hPSC-derived CMs undergo dramatic metabolic changes throughout differentiation^19^. Reduced nicotinamide adenine dinucleotide (phosphate) (NAD(P)H) and oxidized flavin adenine dinucleotide (FAD) are autofluorescent cellular metabolic co-enzymes that can be imaged to collect metabolic information at a single-cell level^20^. The ratio of NAD(P)H to FAD intensity is the “optical redox ratio”, which reflects the relative oxidation-reduction state of the cell. The fluorescence lifetimes of NAD(P)H and FAD are distinct in the free and protein-bound conformations, so changes in these fluorescence lifetimes reflect changes in protein-binding activity^21, 22^. Optical metabolic imaging (OMI) quantifies both NAD(P)H and FAD intensity and lifetime variables. Hence, OMI is suitable to detect the metabolic switches that occur during CM differentiation.

Here, we demonstrate a facile method to non-invasively monitor metabolic changes during hPSC differentiation into CMs by combining OMI with quantitative image analysis. OMI was performed at multiple time points during a 12-day differentiation process under varying conditions (cell density, concentration of Wnt signaling activator) and different hPSC lines (human embryonic pluripotent stem cells and human induced pluripotent stem cells). Differentiation efficiency was quantified by flow cytometry with cTnT labelling on day 12. During the differentiation process all 13 OMI variables, including both NAD(P)H and FAD intensity and lifetime variables, change distinctively between low (< 50% cTnT+ on day 12) and high (≥ 50% cTnT+ on day 12) CM differentiation efficiency conditions. Multivariate analysis found that day 1 cells (24 hours after Wnt activation) formed a distinct cluster from cells at other time points. Logistic regression models based on OMI variables from cells at day 1 performed well for separating low and high differentiation efficiency conditions with a model performance at 0.98 (receiver operating characteristic (ROC) area under the curve (AUC)). This label-free and non-destructive method could be used for quality control for CM manufacturing from hPSCs.

## Results

### NAD(P)H and FAD fluorescence change early in the cardiomyocyte differentiation process

Metabolic state plays an important role in regulating hPSC pluripotency and differentiation^23, 24^, and can be non-invasively monitored via OMI^20, 25^. We recorded the autofluorescence dynamics of NAD(P)H and FAD by OMI during the process of hPSC differentiation into CMs. hPSCs were differentiated following a previous protocol^11^, and cells were imaged on differentiation day 0 (immediately pre-treatment with CHIR99021, a Wnt signaling activator), day 1 (24 hours post-treatment with CHIR99021), day 3 (immediately pre-treatment with IWP2, a Wnt signaling inhibitor), and day 5 (48 hours post-treatment with IWP2). OMI was performed at these time points based on the biphasic role of Wnt signaling activation and inhibition in the CM differentiation protocol^11^. On differentiation day 12, CM differentiation efficiencies were evaluated by flow cytometry with a cardiac specific marker cTnT. Differentiation of CMs from hPSCs critically relies on the timing and the state of Wnt signaling^11^. Both the concentration of CHIR99021^26^ and cell density^7^ are closely related to the activation level of the Wnt signaling pathway. In the current study, CM differentiation efficiencies ranging from nearly 0 to above 60% were achieved by initiating CM differentiation with different CHIR99021 concentrations and hPSC seeding densities (Figure 1a, b, Table 1).

**Figure 1.**
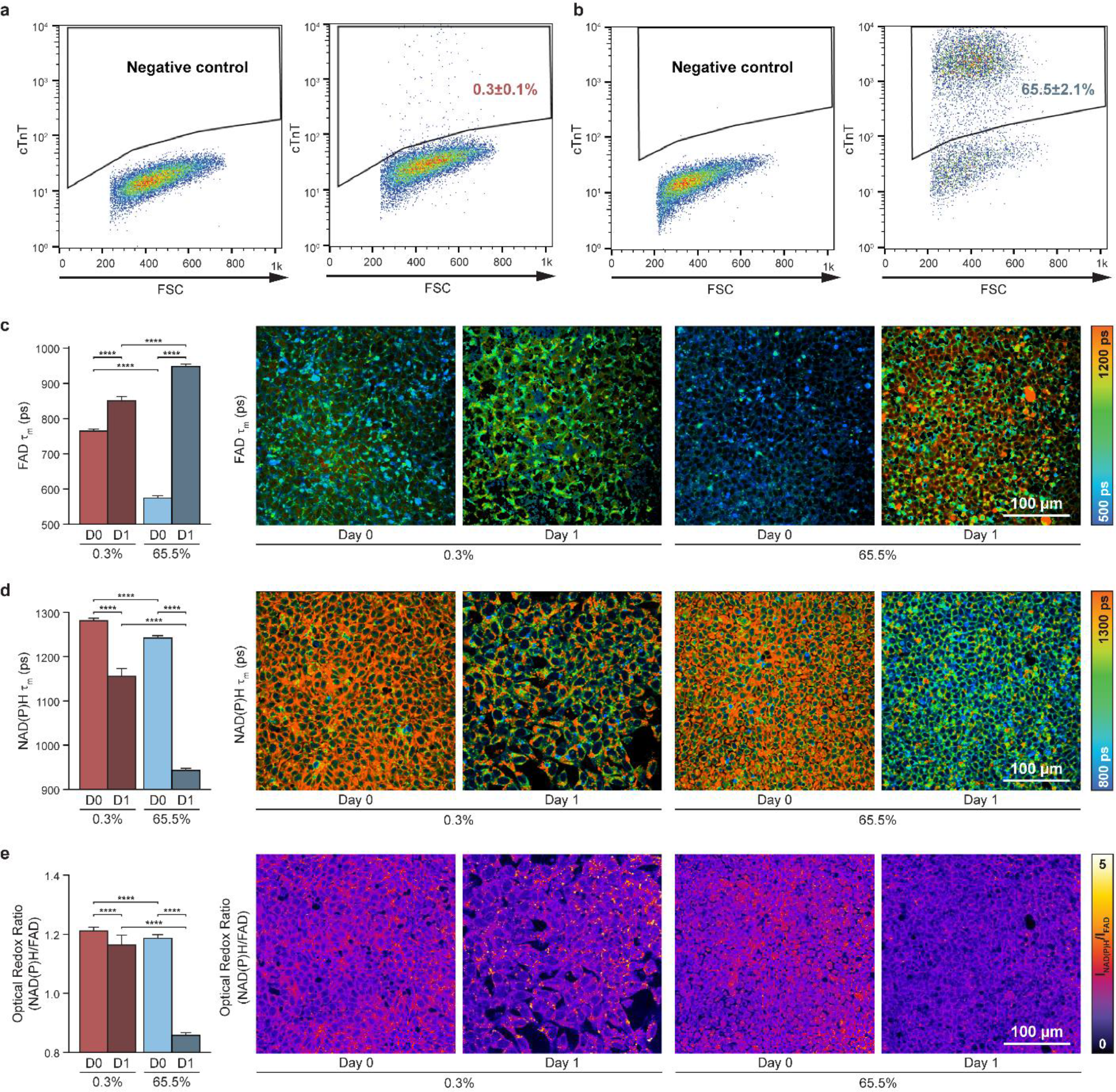
NAD(P)H and FAD fluorescence change differently in the first 24 hours for cells in low vs. high cardiomyocyte differentiation efficiency conditions. hPSCs (H9 embryonic stem cells or IMR90-4 induced pluripotent stem cells) were differentiated into CMs following an established method^11^. On differentiation day 12, cells were verified by flow cytometry with cTnT labelling from three independent replicates. (**a, b**) Representative flow cytometry dot plots for (**a**) low, and (**b**) high differentiation efficiencies along with negative controls. Single-cell quantitative analysis of mean lifetimes (τ_m_, reported as picoseconds) of (**c**) FAD and (**d**) NAD(P)H, and (**e**) optical redox ratio for low differentiation (0.3% cTnT+ on day 12) and high differentiation (65.5% cTnT+ on day 12) efficiencies on day 0 (“D0”, immediately pre-treatment) and day 1 (“D1”, 24 hours post-treatment with CHIR99021, a Wnt signaling activator), and their corresponding representative images. n = 2458, 633, 3534 and 4446 cells for 0.3% day 0, 0.3% day 1, 65.5% day 0, and 65.5% day 1, respectively. Data are presented as mean with 95% CI for each condition each day. Statistical significance was determined by one-way analysis of variance (ANOVA) followed by Tukey’s post hoc tests. ****p< 0.0001. Color bars are indicated on the right.

**Table 1.** Summary of the 11 differentiation conditions. hPSCs, including H9 embryonic stem cells (ESC) or IMR90-4 induced pluripotent stem cells (iPSC) with IMR90 specified, were differentiated into CMs following an established method^11^. On differentiation day 12, cells were verified by flow cytometry with cTnT labelling from three independent replicates to define differentiation efficiency. Data were collected from three biological replicates. Conditions are presented with condition name (seeding density, CHIR99021 concentration, IMR90 status), hPSC line, seeding density, CHIR99021 (Wnt activator) concentration, and differentiation efficiency (mean ± SEM). Low differentiation efficiencies (< 50% cTnT+ on day 12) are shaded in light gray and high differentiation efficiencies (≥ 50% cTnT+ on day 12) are shaded in dark gray.

| Condition | hPSC line | Seeding density (cells/well) | CHIR99021 concentration | Differentiation efficiency |
| --- | --- | --- | --- | --- |
| 1.5m-12μM | H9 ESC | $1.5 \times 10^6$ | 12 μM | $0.3 \pm 0.1\%$ |
| 1.5m-9μM | H9 ESC | $1.5 \times 10^6$ | 9 μM | $0.5 \pm 0.2\%$ |
| 100k-12μM | H9 ESC | $1.0 \times 10^5$ | 12 μM | $0.6 \pm 0.3\%$ |
| 2m-12μM | H9 ESC | $2.0 \times 10^6$ | 12 μM | $15.1 \pm 1.3\%$ |
| 500k-12μM | H9 ESC | $5.0 \times 10^5$ | 12 μM | $19.6 \pm 1.5\%$ |
| 1m-12μM | H9 ESC | $1.0 \times 10^6$ | 12 μM | $21.7 \pm 2.3\%$ |
| IMR90-1m-12μM | IMR90-4 iPSC | $1.0 \times 10^6$ | 12 μM | $26.4 \pm 1.2\%$ |
| IMR90-1m-9μM | IMR90-4 iPSC | $1.0 \times 10^6$ | 9 μM | $38.5 \pm 1.9\%$ |
| 500k-10μM | H9 ESC | $5.0 \times 10^5$ | 10 μM | $51.8 \pm 3.2\%$ |
| IMR90-1m-10μM | IMR90-4 iPSC | $1.0 \times 10^6$ | 10 μM | $53.2 \pm 0.8\%$ |
| 500k-9μM | H9 ESC | $5.0 \times 10^5$ | 9 μM | $65.5 \pm 2.1\%$ |

A total of 13 OMI variables, including the optical redox ratio, NAD(P)H intensity and lifetime variables (τ_1_, τ_2_, α_1_, α_2_, τ_m_), FAD intensity and lifetime variables (τ_1_, τ_1_, α_1_, α_2_, τ_m_) were measured by autofluorescence imaging. The short lifetime (τ_1_) corresponds to free NAD(P)H while the long lifetime (τ_2_) corresponds to protein-bound NAD(P)H. The converse applies to FAD τ_1_ (protein-bound) and τ_2_ (free). Weights are applied to the short (α_1_) and long (α_2_) lifetimes, and the mean lifetime is a weighted average (τ_m_ = α_1_τ_1_ + α_2_τ_2_). Cells under the lowest differentiation efficiency condition (0.3%, Table 1) and highest differentiation efficiency condition (65.5%, Table 1) showed significant differences in OMI variables by day 1. Cells with the highest differentiation efficiency had a lower FAD τ_m_ on day 0 and a higher FAD τ_m_ on day 1 compared to the lowest differentiation efficiency at the same time points (Figure 1c). Similarly, the fold change between day 0 and day 1 for NAD(P)H τ_m_ (Figure 1d) and the optical redox ratio (Figure 1f) is greater for high differentiation efficiency compared to low differentiation efficiency conditions. Significant differences in other OMI variables were observed between day 0 and day 1, as well as between low and high differentiation efficiency conditions (Figure S1). After treating H9 embryonic stem cells with an inhibitor of glycolysis (2-DG)^27^, the optical redox ratio changed oppositely compared to hPSCs undergoing CM differentiation in the first 24 hours (Figure S2a, Figure 1e, Figure S1). However, the optical redox ratio decreased both in H9 embryonic stem cells after rotenone treatment (an oxidative phosphorylation inhibitor)^28^ (Figure S2b) and in hPSCs undergoing CM differentiation in the first 24 hours (Figure 1e). These changes in autofluorescence with known metabolic inhibitors and during CM differentiation indicate that differentiating cells altered their metabolic activity 1 day after CHIR99021 treatment. This observation is consistent with previous studies that found metabolism differed between hPSCs and differentiated cells, and between cells differentiated into CMs and other cell types^29^. Overall, autofluorescence imaging of NAD(P)H and FAD showed significant changes at early time points in the differentiation process, with greater changes in higher CM differentiation efficiency conditions.

### Multivariate analysis reveals unique NAD(P)H and FAD fluorescence in cells 1 day into the differentiation process

To assess differences in OMI variables across days, cells were clustered across all days (day 0, day 1, day 3, and day 5) and differentiation conditions (Table 1) with a Uniform Manifold Approximation and Projection (UMAP) dimension reduction technique^30^. UMAP dimensionality reduction was performed on all 13 OMI variables for projection onto 2D space. UMAP representations of all OMI variables showed a day 1 subpopulation separated from days 0, 3, 5 (Figure 2a, Figure S3a). CM differentiation efficiency conditions were separately evaluated across all days by UMAP. As shown in Figure S3b, day 1 cells (light blue clusters) from high (≥ 50%) differentiation efficiency conditions were distinctly clustered, while cells from low (< 50%) differentiation efficiency conditions clustered together across all days. Therefore, differentiation conditions were separated into low differentiation efficiency (< 50% cTnT+ on day 12, Table 1 shaded in light gray) and high differentiation efficiency (≥ 50% cTnT+ on day 12, Table 1 shaded in dark gray).

**Figure 2.**
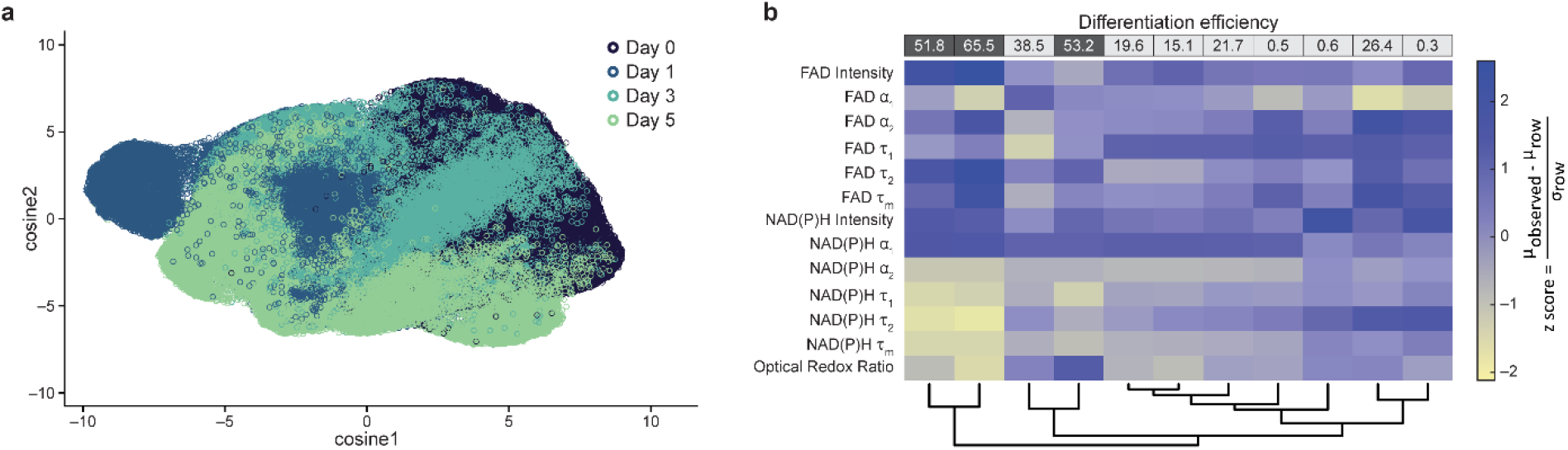
Multivariate analysis reveals unique metabolic profiles in cells differentiated into cardiomyocytes at day 1. **(a)** Uniform Manifold Approximation and Projection (UMAP) dimensionality reduction was performed on all 13 autofluorescence variables (optical redox ratio, NAD(P)H τ_m_, τ_1_, τ_2_, α_1_, α_2_, and intensity; FAD τ_m_, τ_1_, τ_2_, α_1_, α_2_, and intensity) for each cell and projected onto 2D space. Cells from all 11 conditions shown in Table 1 are plotted together. Data include cells from day 0, day 1, day 3, and day 5. Each dot represents one single cell, and n = 25304, 25470, 26228, and 23484 cells for day 0, 1, 3, and 5, respectively. (**b**) Heatmap dendrogram clustering based on similarity of average Euclidean distances across all variable z-scores was performed on day 1 cells across all 11 conditions. Conditions are indicated by the CM differentiation efficiency percentages as noted by column labels at the top of the heatmap (quantified by flow cytometry cTnT+ on day 12, full conditions given in Table 1). Low differentiation efficiencies (< 50%) are shaded in light gray and high differentiation efficiencies are shaded in dark gray (≥ 50%). Z-score = 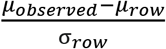, where μ_observed_ is the mean value of together, and σ_row_ is the standard deviation of each variable across all 11 conditions. Autofluorescence variables are indicated on the left side as row labels. n = 25470 cells from day 1.

Heatmap dendrogram clustering based on OMI variable z-scores revealed that cells under high differentiation efficiency conditions on day 1 were clustered closely together and distinct from cells under low differentiation efficiency conditions on day 1 (Figure 2b). Dendrograms of cells on day 0 and day 1 together (Figure S4a) or day 0 alone (Figure S4b) did not show clear separation of high and low differentiation efficiency conditions, indicating that day 1 is the earliest time point to separate low and high differentiation efficiency conditions. In summary, UMAP clustering of all 13 OMI variables across all time points and z-score heatmap clustering from day 0 and day 1 across all differentiation conditions showed that cells under high differentiation efficiency conditions on day 1 clustered together and were distinct from other conditions and time points. Hence, we hypothesize that OMI of live cells on CM differentiation day 1 could predict high and low differentiation efficiencies on day 12.

### OMI variables accurately distinguish cells under low or high differentiation efficiency conditions on day 1

After identifying distinct clustering of day 1 cells in high differentiation efficiency conditions based on all 13 OMI variables, we further explored day 1 OMI data alone. Cells in high differentiation efficiency conditions (Figure 3a, dark gray, ≥ 50% cTnT+ on day 12) formed a distinct cluster from cells under low differentiation efficiency conditions (Figure 3a, light gray, < 50% cTnT+ on day 12) on day 1. However, a small portion of cells from high and low differentiation efficiency conditions overlap. Note that the high differentiation efficiency conditions were not 100% and the low differentiation efficiency conditions were not 0%, so this could explain some of the overlap on day 1.

**Figure 3.**
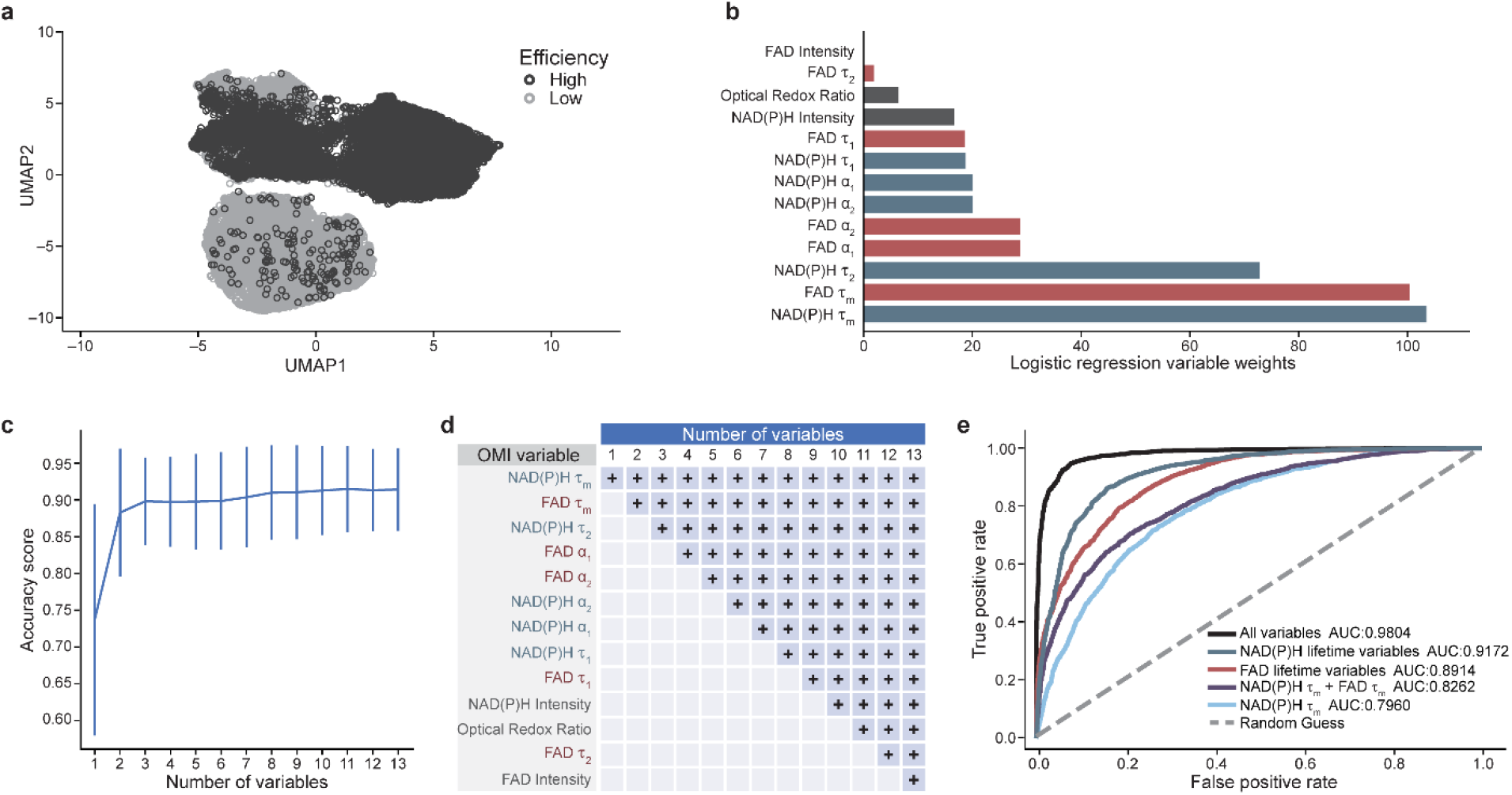
OMI variables accurately distinguish cells under low or high cardiomyocyte differentiation efficiency conditions on day 1. **(a)** UMAP dimensionality reduction was performed on all 13 OMI variables (optical redox ratio, NAD(P)H τ_m_, τ_1_, τ_2_, α_1_, α_2_, and intensity; FAD τ_m_, τ_1_, τ_2_, α_1_, α_2_, and intensity) for each cell on day 1 and projected onto 2D space. Day 1 cells from all 11 conditions shown in Table 1 are plotted together with cells from low (< 50% cTnT+ on day 12) and high (≥ 50% cTnT+ on day 12) CM differentiation efficiencies in light gray and dark gray, respectively. n = 14211 and 11259 cells for low and high differentiation efficiency conditions, respectively. (**b-e**) All OMI data from day 1 cells were randomly partitioned into training and test datasets using 15-fold cross validation, with training and test proportions of 80% and 20%, respectively (n = 20376 cells for training, n = 5094 cells for test). Binary classification was tested for low (< 50% cTnT+ on day 12) vs. high (≥ 50% cTnT+ on day 12) differentiation efficiency conditions on day 1. (**b**) OMI variable weights are shown specific to the logistic regression model. (**c**) Classification accuracy with respect to number of OMI variables was evaluated by chi-squared variable selection to separate low and high differentiation efficiency conditions with the logistic regression model. The number of variables included in the logistic regression model are indicated at bottom-axis. (**d**) The variables included for each logistic regression model [specified by numbers of variables on the x-axis in (**c**)] are defined, where the blue text indicates NAD(P)H lifetime variables and the red text indicates FAD lifetime variables. The OMI variables included in each instance (e.g., 3, 4) are indicated by a light blue + in each column. (**e**) Model performance of the logistic regression classifier was evaluated by receiver operating characteristic (ROC) curves using different OMI variable combinations as labelled. The area under the curve (AUC) is provided for each variable combination as indicated in the legend.

Next, a logistic regression classifier based on all 13 OMI variables was trained on day 1 data to classify cells from low vs. high differentiation efficiency conditions. Variable weights indicated that NAD(P)H τ_m_ and FAD τ_m_ were important variables for discriminating low vs. high differentiation efficiency conditions (Figure 3b). Logistic regression (Figure 3c, d), support vector machine (Figure S5a, b), and random forest (Figure S5c, d) classifiers were generated to test the prediction accuracy using all 13 OMI variables, yielding an accuracy score > 90% for all three classifiers. ROCs based on logistic regression classifiers using all 13 OMI variables and a subset of variables are shown in Figure 3e along with their performance defined by the AUC. Here, the logistic regression classifier using all 13 OMI variables achieves an AUC > 0.98 (Figure 3e). With only NAD(P)H lifetime variables (NAD(P)H τ_m_, τ_2_, α_2_, α_1_, τ_1_) that can be collected in the NAD(P)H channel alone, the AUC is > 0.91 (Figure 3e). The AUC with NAD(P)H lifetime variables is higher than the AUC of other variable subsets, including FAD lifetime variables, NAD(P)H τ_m_ and FAD τ_m_ together, and NAD(P)H τ_m_ alone (Figure 3e). Hence, NAD(P)H lifetime variables alone are sufficient to predict low vs. high CM differentiation efficiency conditions. Additionally, support vector machine and random forest classifiers using all 13 OMI variables achieve an AUC > 0.98 and > 0.99, respectively (Figure S5e). These data indicate that OMI can accurately predict CM differentiation under low vs. high differentiation efficiency conditions at an early time point (day 1).

### Imaging of a cardiac reporter line confirms autofluorescence changes in cells under high differentiation efficiency conditions

Given that OMI can identify CM differentiation efficiency at an early stage, we evaluated a CM reporter line (NKX2.5^EGFP/+^ hPSCs)^31^ to track differentiated CMs together with autofluorescence imaging during the entire differentiation process. The NKX2.5^EGFP/+^ hPSC line expresses EGFP when the cardiac progenitor protein NKX2.5 is expressed, indicating that the cell has differentiated into CM, around differentiation day 7. Although EGFP spectrally overlaps with FAD autofluorescence signals, this interference does not occur until day seven^31^. The final differentiation efficiency was quantified by flow cytometry with cTnT labelling (Figure 4a). Approximately 0.3% and 84.1% CMs were yielded with 12 μM and 9 μM CHIR99021 treatment, respectively.

**Figure 4.**
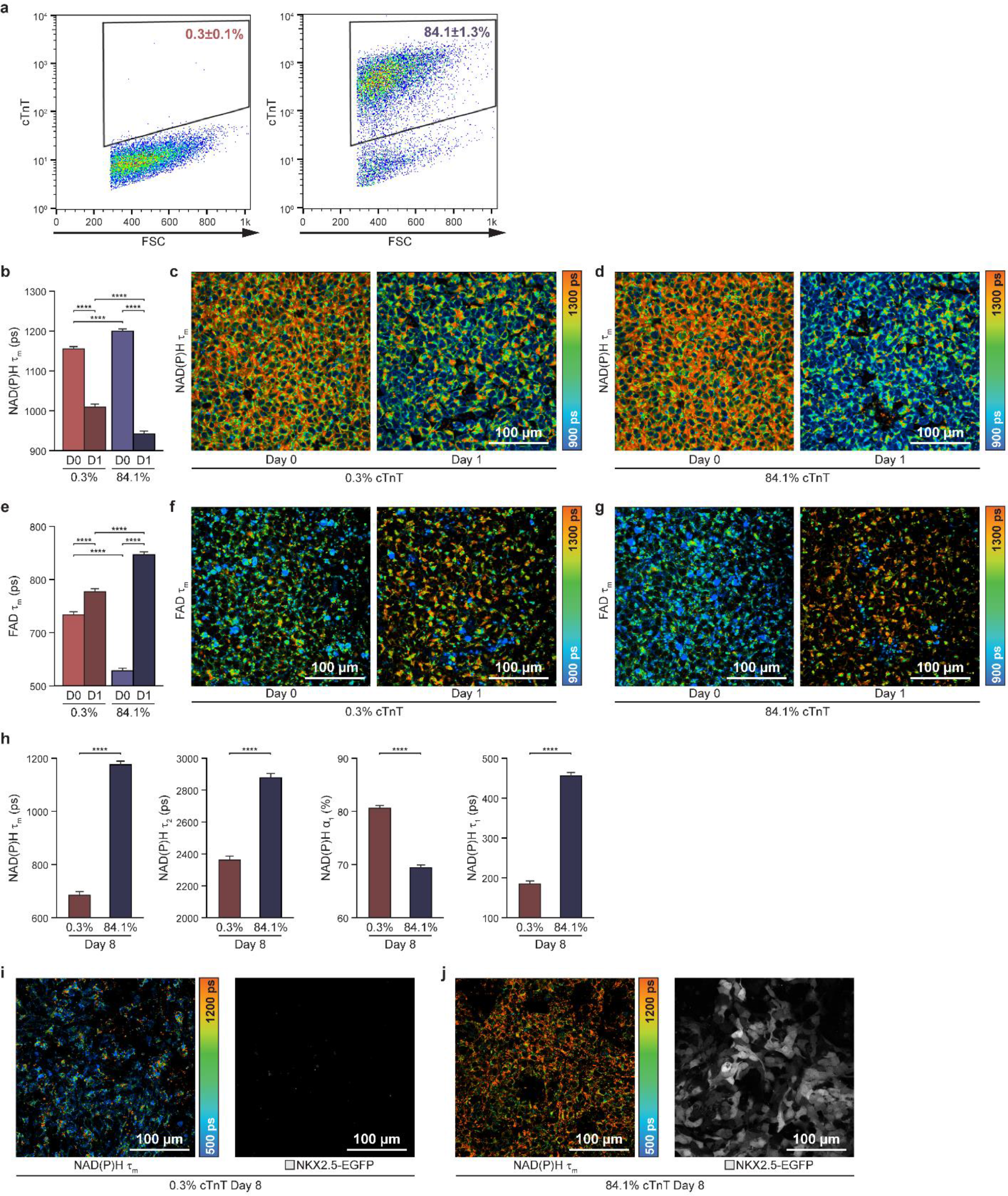
Imaging of a cardiac reporter line confirms autofluorescence changes in cells under high differentiation efficiency conditions. NKX2.5^EGFP/+^ hPSCs were treated with 12 μM and 9 μM of CHIR99021 for the first 24 hours to achieve low and high differentiation efficiencies, respectively. The NKX2.5^EGFP/+^ hPSC line expresses EGFP when the cardiac progenitor protein NKX2.5 is expressed, indicating that the cell has differentiated into CM, around differentiation day 7. (**a)** CM differentiation efficiencies were verified by flow cytometry on day 12 with cTnT labelling. Low differentiation efficiency (12 μM CHIR99021, left) and high differentiation efficiency (9 μM CHIR99021, right) are shown. Data were collected from three biological replicates and presented as mean ± SEM. Single-cell quantitative analysis of (**b**) NAD(P)H mean lifetimes (τ_m_), (**c, d**) representative images, and (**e**) FAD mean lifetimes (τ_m_), (**f, g**) representative images for low (0.3% cTnT+ on day 12) and high (84.1% cTnT+ on day 12) differentiation efficiencies on day 0 (“D0”, immediately pre-treatment) and day 1 (“D1”, 24 hours post-treatment with CHIR99021), respectively. n = 1618, 1017 cells for 0.3% condition day 0; day 1, respectively. n=1633, 1243 cells for 84.1% condition day 0; day 1, respectively. Data are presented as mean with 95% CI. Statistical significance was determined by one-way analysis of variance (ANOVA) followed by Tukey’s post hoc tests. ****p< 0.0001. (**h**) Single-cell quantitative analysis of NAD(P)H τ_m_, τ_2_, α_1_, τ_1_ on differentiation day 8 (Differentiation efficiencies are indicated at the bottom as percent cTnT+ on day 12). Statistical significance was determined by Student’s T-test. **p<0.01. ****p<0.0001. n = 580 and 727 cells for 0.3% and 84.1% condition day 8, respectively. Data are presented as mean with 95% CI. Representative images NAD(P)H τ_m_ and EGFP fluorescence in live cells for (**i**) low differentiation efficiency (0.3% cTnT+) and (**j**) high differentiation efficiency (84.1% cTnT+). ps, picoseconds.

Differences in OMI variables between low (0.3%) and high (84.1%) differentiation efficiencies were assessed with this reporter line. Consistent with the observations in Figure 1c and d, NAD(P)H τ_m_ (Figure 4b-d) and FAD τ_m_ (Figure 4e-g) were significantly different between low vs. high differentiation efficiency conditions on day 0 and day 1 with the CM reporter line. In both conditions, NAD(P)H τ_m_ gradually decreased over the first 5 days of differentiation (Figure S6a). Conversely, FAD τ_m_ gradually increased over the first 5 days for the high differentiation efficiency condition and oscillated over time for the low differentiation efficiency condition (Figure S6b). These observations are consistent with our previous findings (Figure 1, Figure S1) using 11 differentiation conditions across two hPSC lines.

After confirming that EGFP lifetimes do not overlap with NAD(P)H lifetimes (Figure S7), we then evaluated the NAD(P)H lifetimes of differentiated CMs on day 8 when the cells expressed NKX2.5-EGFP. As shown in Figure 4h-j, NAD(P)H lifetime variables (τ_m_, τ_2_, α_1_, τ_1_) were significantly different between low and high differentiation efficiencies on day 8. Similar changes in NAD(P)H lifetime variables were also observed in H9 embryonic stem cells treated with an inhibitor of glycolysis (2-DG)^27^ (Figure S8). In summary, with a cardiac reporter line, we further confirmed that NAD(P)H and FAD fluorescence variables reflect CM differentiation efficiency from hPSCs. Differentiated CMs (84.1%) exhibit dramatically different autofluorescence compared to differentiated non-CMs (0.3%), which provides further evidence that OMI can discriminate between CMs and non-CMs after the differentiation process is complete.

## Discussion

Here, we report a non-invasive label-free imaging method to predict the outcome of hPSC differentiation into CMs. By combining live cell autofluorescence lifetime imaging, single-cell image analysis, and machine learning, we robustly separate low (< 50%) from high (≥ 50%) CM differentiation efficiency conditions as early as day 1. The prediction accuracy was over 90% and the model performance was 0.98 (AUC of ROC) with all 13 OMI variables combined across 11 different differentiation conditions including 3 hPSC lines.

Recent evidence links Wnt signaling and glycolytic activities during hPSC differentiation into mesoderm ^32, 33^. Consistent with these findings, changes in OMI variables on day 1 of CM differentiation (24 hours after Wnt activation, Figure 1) and with known metabolic inhibitors in stem cells (Figure S2, Figure S8) indicate that changes in OMI variables are due to increased glycolytic activity on day 1 of CM differentiation. Considering the important role of Wnt signaling activation in mesoderm and CM differentiation^11^, and embryonic development^34^, greater changes in OMI variables in the high differentiation efficiency condition could indicate more glycolytic activity due to successful Wnt activation compared to the low differentiation efficiency condition (Figure 1c-e, Figure S1). Taken together, these results reveal that autofluorescence imaging can separate CM differentiation efficiencies at an early stage based on metabolic changes (Figure 3b-e).

At the end of our differentiation process, some cells were not positive for cTnT and therefore were not CMs. Previous studies have shown that these non-CMs at the end of the differentiation process are primary cardiac-like fibroblasts together with a small portion of non-differentiated hPSCs^35^. hPSC-derived CMs exhibit distinct metabolism from other hPSC-derived non-CMs^36^. Co-culture of cardiac fibroblasts and CMs can induce fibroblast glycolytic activity and lactate secretion from fibroblasts^37^. Co-culture of cardiac fibroblasts and CMs also promotes a more mature phenotype in CMs along with increased reliance on oxidative phosphorylation^35^. Similarly, differences in NAD(P)H lifetime variables between CMs and non-CMs on day 8 (Figure 4h) are consistent with decreased glycolytic activity in the CMs (Figure S8).These results further confirm that autofluorescence imaging can distinguish the distinct metabolic activities between hPSC-derived CMs and other non-CMs.

We have demonstrated that autofluorescence imaging can resolve metabolic changes in CM differentiation and predict the differentiation outcome at early time points. However, our method has limitations. The differentiation efficiency of hPSCs is susceptible to cell line variability, cell culture microenvironment, and differentiation protocol^38^. We note that the differentiation efficiency measured from flow cytometry in our experiments was not higher than 90%. This may be due to photo-toxicity during the imaging process that may moderately interrupt CM differentiation. In future studies, good manufacturing practice standards could be applied to optimize the evaluation process and minimize the interruption on differentiation. Additionally, OMI relies on only two metabolites, NAD(P)H and FAD, that do not comprehensively characterize cellular metabolic activities. More mechanistic studies together with other assays, including metabolite liquid chromatography–mass spectrometry^23^, NMR spectrometry^39^, single-cell RNASeq^40^, and quantitative proteomics^41^, need to be performed to reveal the relationship between metabolic dynamics and hPSC differentiation into CMs. Additionally, alternative differentiation protocols will require algorithms trained on OMI data in these new conditions to robustly classify differentiation efficiencies.

Overall, we developed a non-invasive method to predict the efficiency of hPSC differentiation into CMs at early differentiation stages. hPSCs hold great promise for regenerative medicine and pharmaceutical development, but large-scale cell manufacturing suffers from variability across hPSC lines and cell culture conditions. Our studies indicate that autofluorescence can predict CM differentiation efficiency at an early stage, which could enable real-time and/or in-line monitoring during cell manufacturing. This method could lower manufacturing costs and personnel time by flagging samples for timely interventions. Similar technologies could also impact other areas of regenerative cell manufacturing.

## Materials and methods

### hPSC culture and cardiomyocyte differentiation

Human H9 embryonic stem cells^42^, human IMR90-4 induced pluripotent stem cells^43^, and NKX2.5^EGFP/+^ hPSCs31 were maintained on Matrigel (Corning)-coated surfaces in mTeSR1 (STEMCELL Technologies) as previously described^44^. CM differentiation was performed as described previously^11^. Different cell seeding densities and different concentrations of CHIR99021 were applied to manipulate differentiation efficiency. Briefly, hPSCs were singularized with Accutase (Thermo Fisher Scientific) and plated onto Matrigel-coated plates at a density ranging from of 2.9 × 10^4^ cells/cm^2^ to 5.7 × 10^5^ cells/cm^2^ (1.0 × 10^5^ cells to 2.0 × 10^6^ cells per well of a 12-well plate) in mTeSR1 supplemented with 10 μM Rho-associated protein kinase (ROCK) inhibitor Y-27632 (Selleckchem) 2 days before initiating differentiation. Differentiation was initiated by Wnt signaling activation with 8 μM to 12 μM CHIR99021 (Selleckchem) on day 0, followed by inhibition of Wnt signaling with 5μM IPW2 on day 3.

### Flow cytometry

Cells on differentiation day 12 were disassociated with Accutase, fixed in 1% PFA for 15 minutes at room temperature, and then blocked with 0.5% bovine serum albumin (BSA) with 0.1% Triton X-100. Cells were then stained with primary antibody anti-cTnT (Lab Vision; 1:200) and secondary antibody (Thermo Fisher; goat anti-mouse, Alexa Fluor 488; 1:500) in 0.5% BSA with 0.1% Triton X-100. Data were collected on a FACSCalibur flow cytometer and analyzed with FlowJo. Data were collected from three biological replicates and presented as means ± SEM. cTnT positive percentage was rounded up at one decimal place.

### Autofluorescence Imaging of NAD(P)H and FAD

Fluorescence lifetime imaging (FLIM) was performed by an Ultima two-photon imaging system (Bruker) composed of an ultrafast tunable laser source (Insight DS+, Spectra Physics) coupled to a Nikon Ti-E inverted microscope with time-correlated single photon counting electronics (SPC-150, Becker & Hickl, Berlin, Germany). The ultrafast tunable laser source enables sequential excitation of NAD(P)H at 750 nm and FAD at 890 nm. NAD(P)H and FAD emission was isolated using 440/80 nm and 550/100 nm bandpass filters (Chroma), respectively. The laser power at the sample for NAD(P)H and FAD excitation was approximately 2.3 mW and 7.9 mW, respectively. Fluorescence lifetime decays with 256 time bins were acquired across 256 × 256 pixel images with a pixel dwell time of 4.8 μs and an integration period of 60 seconds. All samples were illuminated through a 40×/1.15 NA objective (Nikon). FLIM was performed on differentiation day 0 (immediately pre-treatment with CHIR99021, a Wnt signaling activator), day 1 (24 hours post-treatment with CHIR99021), day 3 (immediately pre-treatment with IWP2, a Wnt signaling inhibitor), and day 5 (48 hours post-treatment with IWP2). For NKX2.5^EGFP/+^ hPSCs, day 8 NAD(P)H lifetime variables were also collected. For the 2DG experiment, H9 embryonic stem cells were imaged before and 2 hours after 10 mM 2DG treatment, respectively. For the rotenone experiment, H9 embryonic stem cells were imaged before and 15 minutes after 10 μM rotenone treatment, respectively.

### Image analysis

Lifetime images of NAD(P)H and FAD were analyzed via SPCImage software (Becker & Hickl). Two-component decays were calculated by the following equation^22^: 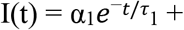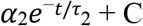. Fluorescence intensity images were generated by integrating photon counts over the per-pixel fluorescence decays. The per-pixel ratio of NAD(P)H fluorescence intensity to FAD intensity was calculated to determine optical redox ratio. A customized CellProfiler pipeline was used to segment individual cell cytoplasms^45^. Cytoplasm masks were applied to all images to determine single-cell optical redox ratio and NAD(P)H and FAD fluorescence lifetime variables. Fluorescence lifetime variables consist of the mean lifetime (τ_m_ = α_1_τ_1_ + α_2_τ_2_), free- and protein-bound lifetime components (τ_1_ and τ_2_ for NAD(P)H, and τ_2_ and τ_1_ for FAD, respectively), and their fractional contributions (α_1_ and α_2_; where α_1_ + α_2_ = 1) for each individual cell cytoplasm. A total 13 OMI variables were analyzed for each cell cytoplasm: FAD intensity, FAD α_1_, FAD α_2_, FAD τ_1_, FAD τ_2_, and FAD τ_m_; NAD(P)H intensity, NAD(P)H α_1_, NAD(P)H α_2_, NAD(P)H τ_1_, NAD(P)H τ_2_, and NAD(P)H τ_m_; optical redox ratio= 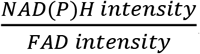.

The phasor plot of lifetime decays for enhanced green fluorescent protein (EGFP) and NAD(P)H was performed as previously described^46^. Briefly, phasor lifetime plots are derived from a Fourier transformation of fluorescence lifetime decay curves by a custom algorithm. The fluorescence lifetime of each pixel in the image is presented in a 2D phasor plot with the unitless horizontal axis (G) and the vertical axis (S).

### UMAP clustering

Clustering of cells across all days and differentiation efficiency conditions was represented using Uniform Manifold Approximation and Projection (UMAP). UMAP dimensionality reduction was performed on all 13 OMI variables (optical redox ratio, NAD(P)H τ_m_, τ_1_, τ_2_, α_1_, α_2_, and intensity; FAD τ_m_, τ_1_, τ_2_, α_1_, α_2_, and intensity) for projection in 2D space. The following parameters were used for UMAP visualizations: “n _neighbors”: 10; “min_dist”: 0.3, “metric”: cosine, “n_components”: 2.

### Classification methods

Logistic regression classifiers were trained to distinguish cells at low (< 50% cTnT+ on day 12) and high (≥ 50% cTnT+ on day 12) differentiation efficiency 1 day post-treatment with CHIR99021. Consistent separation of day 1 UMAP clusters from all other days across differentiation conditions prompted classification of single-cell autofluorescence data from day 1 differentiation. All day 1 OMI data were randomly partitioned into training and test datasets using 15-fold cross validation for training and test proportions of 80% and 20%, respectively (n = 20376 cells in the training set, n = 5094 cells in the test set). Chi-squared variable selection was used to evaluate classification accuracy as a function of the number of training variables. Variable weights for OMI variables were extracted to determine the contribution of each variable to the trained logistic regression model. Receiver operating characteristic (ROC) curves were generated to evaluate the logistic regression model performance on classification of test set data. Support vector machine and random forest classifiers were also trained to classify low and high differentiation efficiencies on day 1 to determine whether classification performance was dependent on the chosen model. Training and test set partitioning and variable selection methods for support vector machine and random forest classifiers were identical to those reported for the logistic regression model.

### Z-score hierarchical clustering

Z-score of each OMI variable for each condition was calculated across all 11 conditions. Z-score = 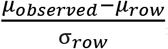, where μ_observed_ is the mean value of each variable for each condition; μ_row_ is the mean value of each variable for all 11 conditions together, and σ_row_ is the standard deviation of each variable across all 11 conditions. Heatmaps of z-scores for all OMI variables were generated to visualize differences in each variable between low and high differentiation efficiency conditions at day 0 and day 1. Dendrograms show clustering based on similarity of average Euclidean distances across all variable z-scores. Heatmaps and associated dendrograms were generated in R (heatmap.2, gplots package).

### Statistics

Data for OMI variables are presented as mean with 95% CI. Data for flow cytometry are presented as mean ± SEM. Statistical significance was determined by Student’s T-test (two-tailed) between two groups. Three or more groups were analyzed by one-way analysis of variance (ANOVA) followed by Tukey’s post hoc tests. P < 0.05 was considered statistically significant and indicated in the figures.

## Author contributions

The project was conceived by T.Q., and M.C.S. The experiments were designed by T.Q. and M.C.S, and were carried out by T.Q., Data analysis was performed by T.Q., T.M.H., A.R.H., K.S., and K. S. The manuscript was written by T.Q., T.M.H., and M.C.S.

## Data and materials availability

All data needed to evaluate the conclusions in the paper are present in the paper and/or the Supplementary Materials. Additional data related to this paper may be requested from the authors.

## Supplemental figures and figure legends

**Figure S1.**
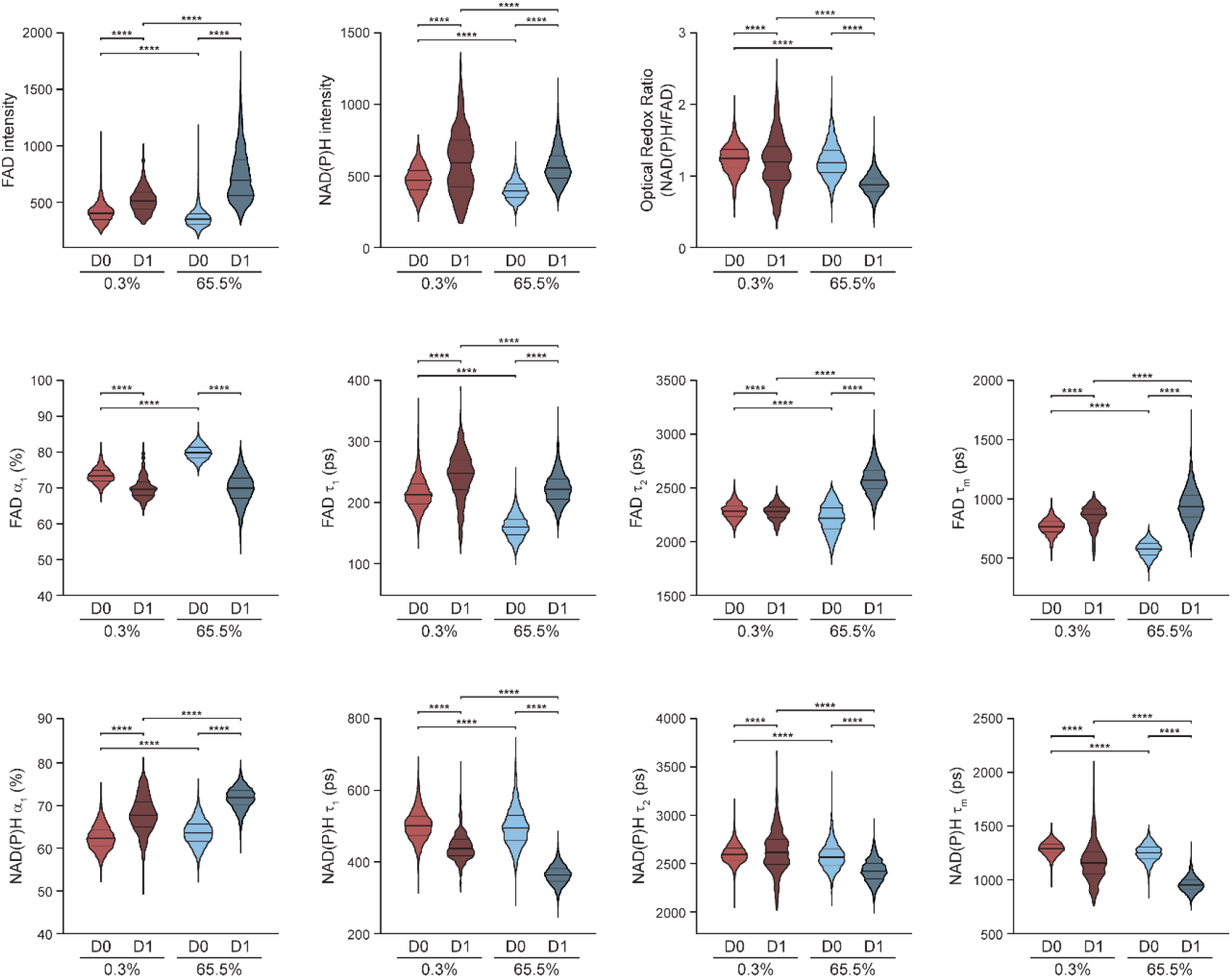
NAD(P)H and FAD fluorescence change differently in the first 24 hours for cells in low vs. high cardiomyocyte differentiation efficiency conditions. hPSCs were differentiated into CMs following an established method^11^. Single-cell quantitative analysis of OMI variables, including FAD intensity, NAD(P)H intensity, optical redox ratio, FAD α_1_, τ_1_, τ_2_, τ_m_, and NAD(P)H α_1_, τ_1_,τ_2_, τ_m_, for low differentiation (0.3% cTnT+ on day 12) and high differentiation (65.5% cTnT+ on day 12) efficiencies on day 0 (“D0”, immediately pre-treatment) and day 1 (“D1”, 24 hours post-treatment with CHIR99021). n = 2458, 633, 3,534, and 4446 cells for 0.3% D0, 0.3% D1, 65.5% D0, and 65.5% D1, respectively. On differentiation day 12, cells were verified by flow cytometry with cTnT labelling. Data are presented in violin plots with middle bar for the mean, lower bar for 25 percentiles, and upper bar for 75 percentiles. Statistical significance was determined by one-way analysis of variance (ANOVA) followed by Tukey’s post hoc tests. ****p< 0.0001. ps, picoseconds.

**Figure S2.**
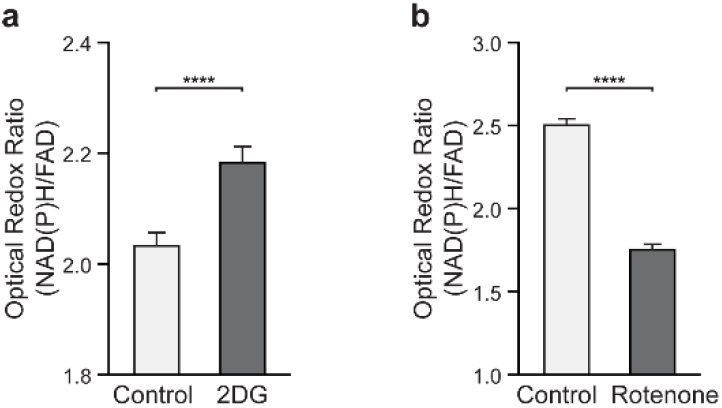
Changes of optical redox ratio after treatment with 2DG or rotenone. (**a**) Single-cell quantitative analysis of optical redox ratio for H9 embryonic stem cells before and 2 hours after 10 mM 2DG treatment. n = 1051 and 900 cells for before and after 2DG treatment, respectively. (**b**) Single-cell quantitative analysis of optical redox ratio for H9 embryonic stem cells before and 15 minutes after 10 μM rotenone treatment. n = 1042 and 986 cells for before and after rotenone treatment, respectively. Data are presented as mean with 95% CI. Statistical significance was determined by Student’s T-test. ****p<0.0001. ps, picoseconds.

**Figure S3.**
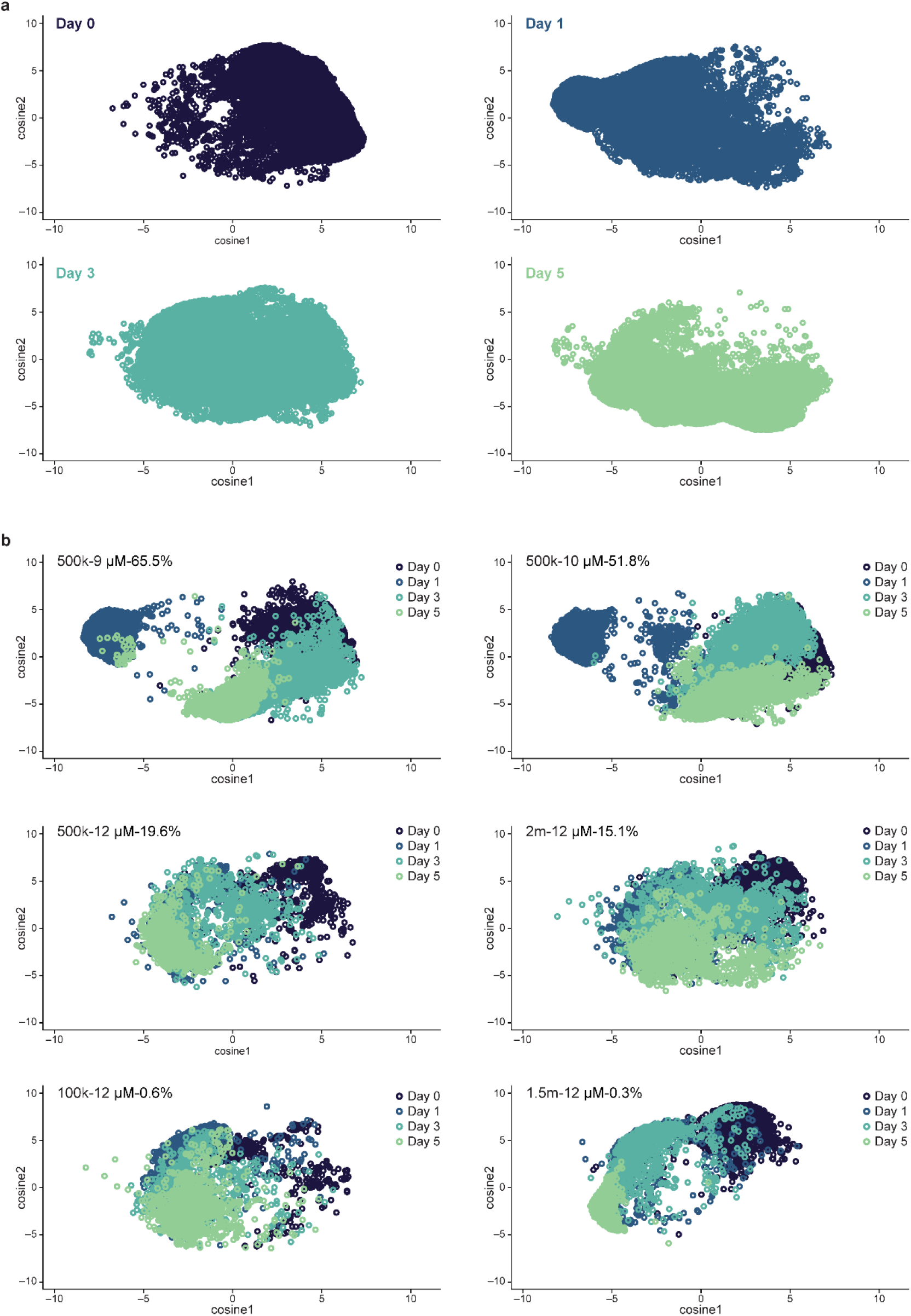
UMAP plotted by separate days and conditions. **(a)** Uniform Manifold Approximation and Projection (UMAP) dimensionality reduction was performed on all 13 autofluorescence variables (optical redox ratio, NAD(P)H τ_m_, τ_1_, τ_2_, α_1_, α_2_, and intensity; FAD τ_m_, τ_1_, τ_2_, α_1_, α_2_, and intensity) for each cell and projected onto 2D space. Cells from all 11 conditions shown in Table 1 are plotted by days separately. Each dot represents one single cell, and n = 25304, 25470, 26228, and 23484 cells for day 0, 1, 3, and 5, respectively. (**b**) Separated UMAP clusters for each differentiation condition. Conditions are labelled with original cell seeding density, CHIR99021 treatment concentration, and final cardiomyocyte differentiation efficiency quantified by flow cytometry (detailed in Table 1). n = 13897, 13852, 4357, 7601, 3445, and 5526 cells for condition 65.5%, 51.8%, 19.6%, 15.1%, 0.6%, and 0.3%, respectively.

**Figure S4.**
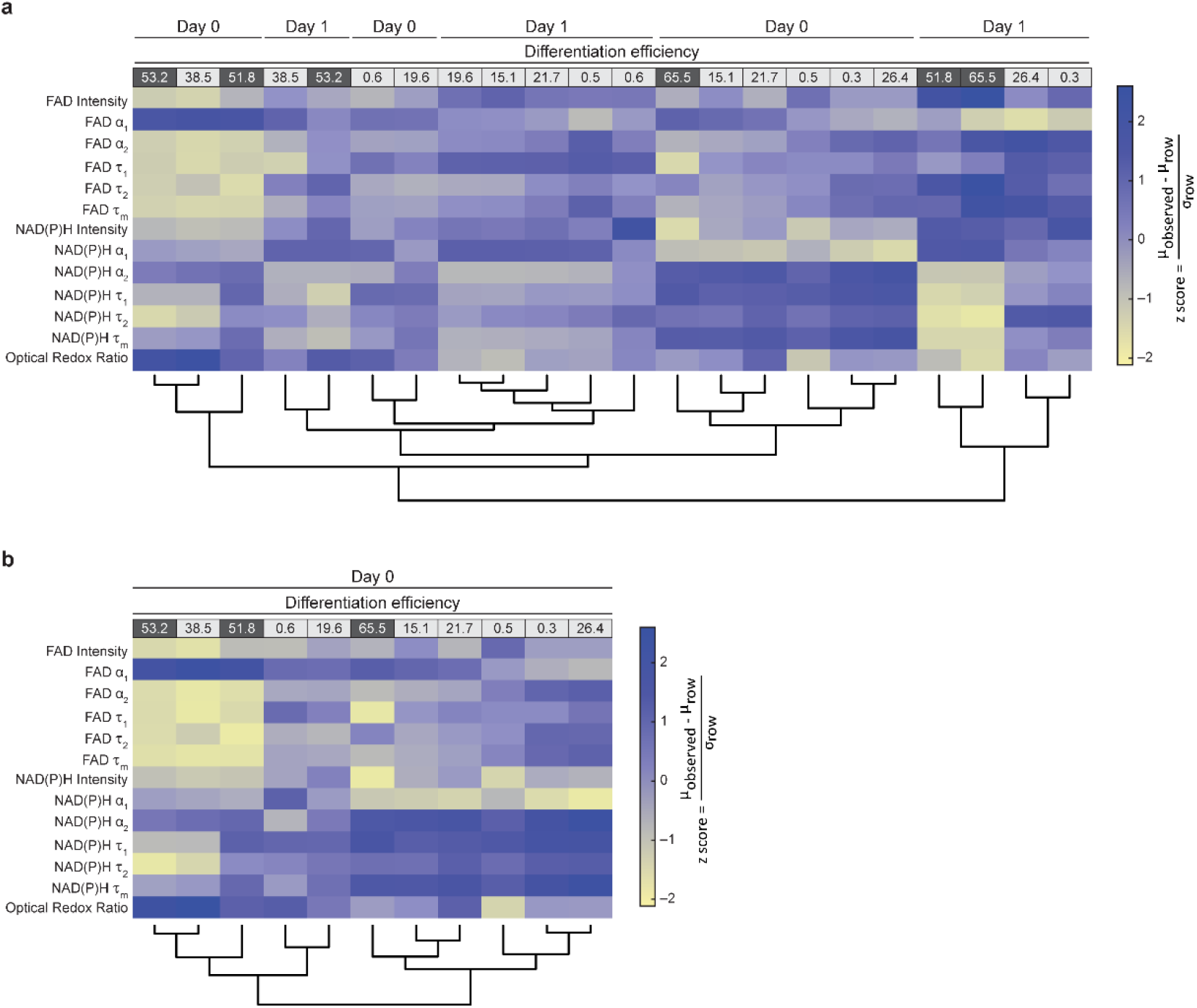
Heatmap dendrogram clustering of OMI variable z-score on day 0 and day 1. Heatmap dendrogram clustering based on similarity of average Euclidean distances across all OMI variable z-scores was performed on (**a**) day 0 (immediately pre-treatment) and day 1 cells (24 hours post-treatment with CHIR99021) across all 11 conditions together or (**b**) day 0 alone. Conditions are indicated by the CM differentiation efficiency percentages as noted by column labels at the top of the heatmap (quantified by flow cytometry cTnT+ on day 12, full conditions given in Table 1). Low differentiation efficiencies (< 50%) are shaded in light gray and high differentiation efficiencies are shaded in dark gray (≥ 50%). Z-score = 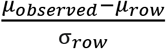, where μ_observed_ is the mean value of each variable for each condition, μ_row_ is the mean value of each variable for all 11 conditions together, and σrow is the standard deviation of each variable across all 11 conditions. Autofluorescence variables are indicated on the left side as row labels. n = 25304 and 25470 cells for day 0, 1, respectively.

**Figure S5.**
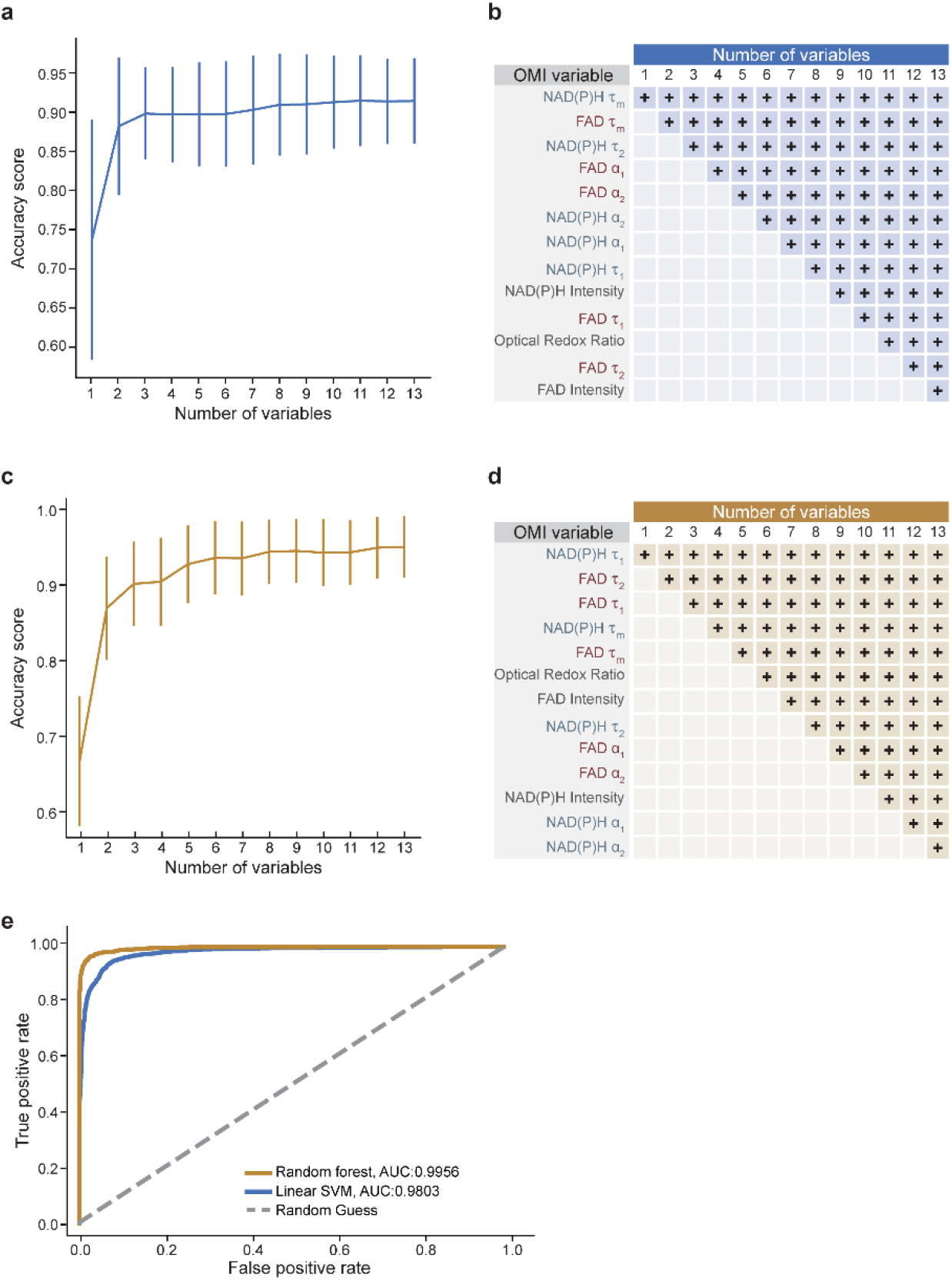
Support vector machine and random forest models for classification of cells on day 1. All OMI data (optical redox ratio, NAD(P)H τ_m_, τ_1_, τ_2_, α_1_, α_2_, and intensity; FAD τ_m_, τ_1_, τ_2_, α_1_, α_2_, and intensity) from day 1 cells were randomly partitioned into training and test datasets using 15-fold cross validation for training and test proportions of 80% and 20%, respectively (n = 20376 cells for training, 5094 cells for test). Classification accuracy with respect to number of OMI variables was evaluated by chi-squared variable selection to separate low (< 50% cTnT+ on day 12) and high (≥ 50% cTnT+ on day 12) differentiation efficiency conditions by (**a**) support vector machine (SVM). (**b**) The variables included for each SVM [specified by numbers of variables on the x-axis in (**a**)]. The OMI variables included in each instance (e.g., 3, 4) are indicated by a blue + in each column. (**c**) Classification accuracy with respect to number of OMI variables for random forest classification. (**d**) The variables included for each random forest classifier [specified by numbers of variables on the x-axis in (**c**)]. The OMI variables included in each instance are indicated by a yellow + in each column (**e**) Model performance was evaluated by receiver operating characteristic (ROC) curves displaying classification performance with the SVM (blue curve) or random forest model (yellow curve) using all 13 OMI variables. The area under the curve (AUC) is provided for SVM and random forest classifiers as indicated in the legend.

**Figure S6.**
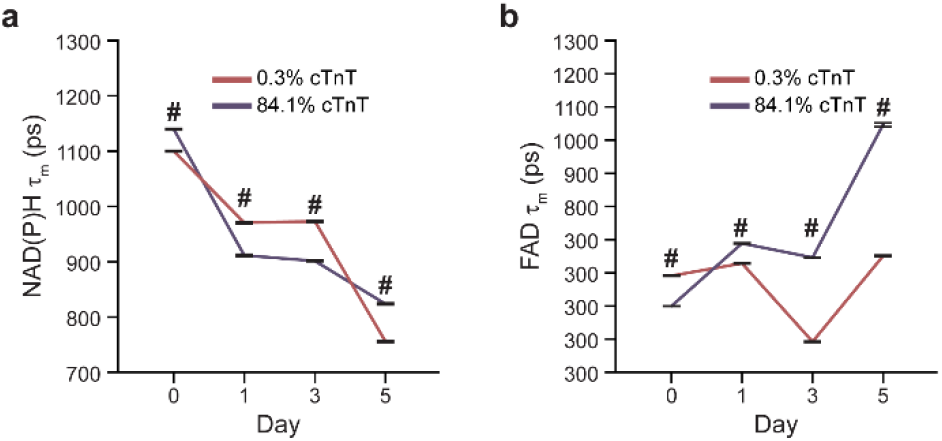
Changes in lifetimes of NAD(P)H and FAD during the first 5-days of differentiation. Single-cell quantitative analysis of mean lifetimes (τ_m_) of (**a**) NAD(P)H and (**b**) FAD on days 0, 1, 3, and 5. Paired student T-test was performed between low (0.3% cTnT+) and high (84.1% cTnT+) differentiation efficiency conditions on each day. #p<0.0001 for 0.3% vs. 84.1% differentiation efficiency conditions on day 0, day 1, day 3, and day 5, respectively. Data were collected from over 500 cells for each day for each condition and presented as mean ± SEM. ps, picoseconds.

**Figure S7.**
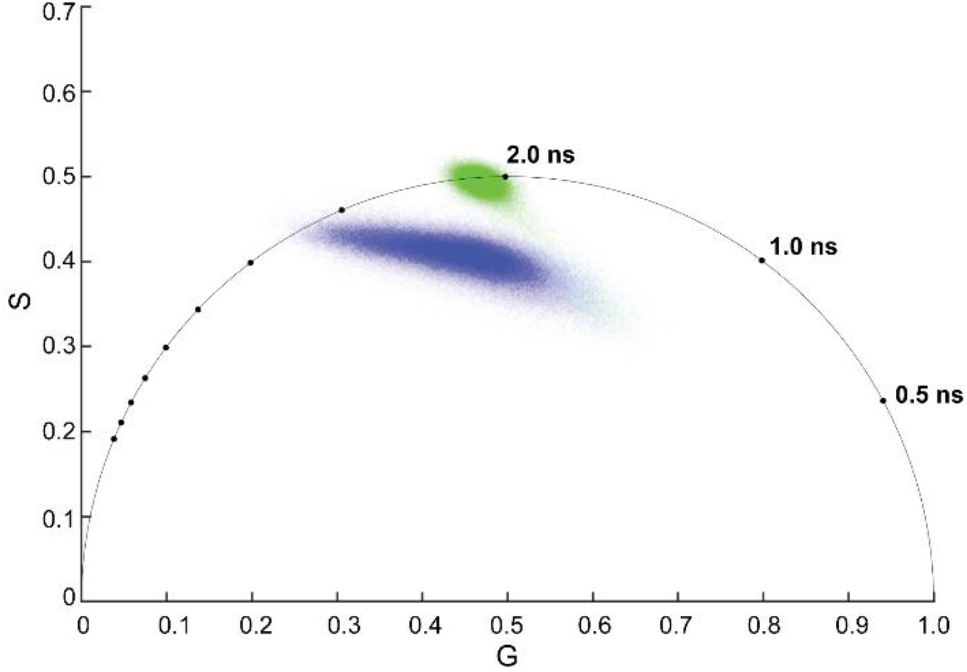
Representative phasor plots reveal separation of NAD(P)H and EGFP decays. Phasor plots of fluorescence decays provide a visual distribution of the molecular species in an image by clustering pixels with similar lifetimes, which allows assessment of overlap between lifetime species. The phasor fluorescence decay plots are derived from a Fourier transformation of the fluorescence lifetime decay^46^. The fluorescence lifetime of each pixel in the image is presented in the 2D phasor plot with the unitless horizontal (G) and vertical (S) axes. Blue dots are NAD(P)H decays and green dots are EGFP decays. The lifetimes of 3 positions on the unit circle are given in nanoseconds for reference.

**Figure S8.**
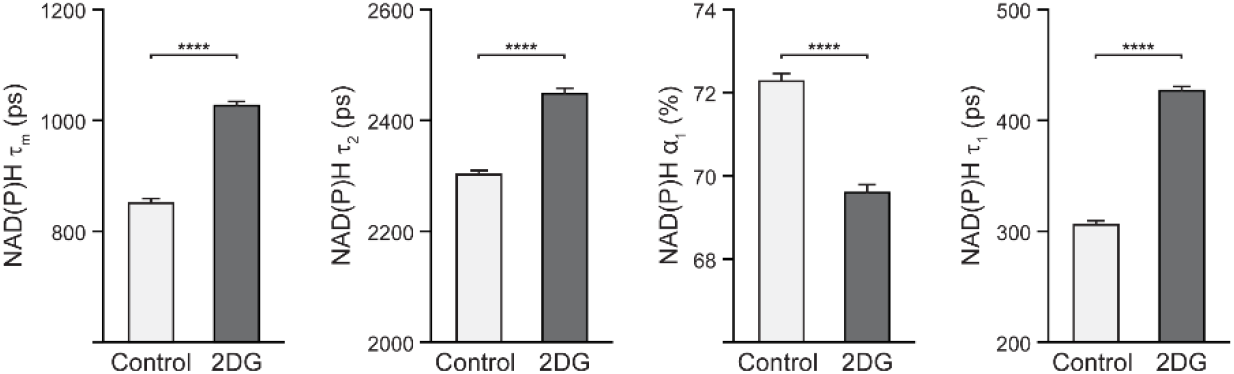
Changes of NAD(P)H lifetime variables after treatment with 2DG. Single-cell quantitative analysis of NAD(P)H lifetime variables (τ_m_, τ_2_, α_1_, τ_1_) for H9 embryonic stem cells before and 2 hours after 10 mM 2DG treatment. n = 1051 and 900 cells for before and after 2DG treatment, respectively. Data are presented as mean with 95% CI. Statistical significance was determined by Student’s T-test. ****p<0.0001. ps, picoseconds.

